# Dynamic Brain Network States during Suspenseful Film Viewing

**DOI:** 10.1101/2021.10.19.465030

**Authors:** Matthew A. Bezdek, Shella D. Keilholz, Eric H. Schumacher

## Abstract

Studies of brain activity evoked by naturalistic stimuli have found narrative suspense produces reliable patterns of activation consistent with increased attention to the narrative and decreased processing of non-narrative stimuli (Bezdek et al., 2015). Yet the progression of brain states (i.e. patterns of activity levels across the whole brain) at suspense peaks and how they differ from suspense valleys or resting state scans has not previously been investigated. We used a fast fMRI acquisition sequence to measure sub-second brain states at peaks and valleys of narrative suspense and during rest. We found five brain states that occur during suspenseful film viewing, four of which differ in their frequency near suspense peaks compared to valleys. The pattern of results is consistent with the theoretical view that suspense captures and focuses attentional processing, triggering subsequent cognitive processing of narrative events.

---

Activity in networks across the brain changes dynamically over the course of both resting state and task-based scans. While the patterns of brain activation over time vary across participants during resting state scans, during the naturalistic presentation of edited commercial films, responses can be highly synchronized across sets of participants, as measured with the locations of eye fixations (Mital et al., 2011; Wang et al., 2012), electroencephalography (EEG; Dmochowski et al., 2012), and functional magnetic resonance imaging (fMRI; Hasson et al., 2004, 2010). Suspenseful film stimuli in particular have been shown to elicit time-locked patterns of behavioral and neural activity across participants (Bezdek et al., 2015, 2017; Bezdek & Gerrig, 2017; Dmochowski et al., 2012; Hasson et al., 2008).

Prior research using univariate BOLD analyses has identified networks of brain regions that reliably increase or decrease in activity as a function of narrative suspense. Moments of increasing narrative suspense produce increased activity in higher-order visual processing regions and the ventral attention network and decreased activity in the default mode network (Bezdek et al., 2015, 2017). These findings complement behavioral studies of suspenseful films that have reported suspense impairs performance on secondary tasks (Bezdek & Gerrig, 2017; Cohen et al., 2015).

Yet the slow acquisition of traditional whole-brain fMRI recording has limited the sensitivity to study dynamic changes in brain activity evoked by changes in suspense. Using a multiband sequence to excite several slices simultaneously can boost the spatial or temporal resolution of fMRI data. Previous studies have reported that using multiband sequences to acquire faster observations can boost statistical power over the noise inherent in fMRI acquisition (Boyacioğlu et al., 2015; Todd et al., 2016), though multiband sequences have also been reported to produce artifacts on the recorded images, particularly at higher acceleration factors (Demetriou et al., 2018; Preibisch et al., 2015).

Because multi-band sequences can increase the temporal resolution of fMRI, they may be more effective than conventional sequences at measuring brain states as they change over the duration of an fMRI run. By ‘brain state’, we mean a pattern of the static spatial distribution of activation and deactivation across the entire brain at a particular time. A brain may traverse between brain states according to endogenous and exogenous task demands during the flow of experience (Bezdek et al., 2019). One method for classifying brain states over time is to apply a clustering algorithm to group time points (individual volumes) that are maximally similar (Chen et al., 2015; Liu & Duyn, 2013; Smith et al., 2018). This method of measuring dynamic whole-brain states provides more sensitive temporal resolution than alternative methods like sliding window analyses, which require balancing the competing demands of choosing a short window to increase temporal specificity with choosing a long enough window to generate reliable correlation estimates (Hutchison et al., 2013). Prior research using clustering of time points to classify brain states has identified reliable states across a working memory task and rest (Chen et al., 2015), and across sessions of resting state scans (Smith et al., 2018). However, this method has not previously been used to identify states during the viewing of naturalistic suspenseful films.

In the present study, we used a fast multiband fMRI sequence to measure brain activity during suspenseful film excerpts. The occurrence of discrete brain states was compared between peaks and valleys in the time course of narrative suspense. The composition of brain states and the transitions between them was also compared between movie watching and rest.

## Method

### Participants

Thirteen participants (8 female, age range 18-25 years) were included in the fMRI analyses. Participants were screened to ensure they were right-handed and did not have any implants or neurological conditions that would interfere with fMRI safety or imaging quality. An additional two participants began participation in the study but did not complete the fMRI session due to technical difficulties or discomfort in the scanner. Of the 13 participants included in analyses, one had excessive head movement during the resting scan and another one had excessive head movement during the movie runs. After excluding these scans, there were 12 participants included in the movie and 12 participants in the resting state analyses, with one participant in each analysis who did not appear in the other analysis. Participants were compensated with $10 per hour for their participation. All procedures were approved by the Institutional Review Board of the Georgia Institute of Technology.

### Stimuli

Excerpts from ten feature films were used as stimuli (*Alien, Blood Simple, Cliffhanger, Léon: The Professional, License to Kill, The Man Who Knew Too Much, Marnie, Misery, Munich*, and *North by Northwest*). These excerpts have been used in previous research to elicit dynamic feelings of suspense (Bezdek et al., 2015, 2017; Bezdek & Gerrig, 2017). In each excerpt, a character faces the threat of an extremely negative outcome. Over the course of each excerpt, pieces of new narrative information make the potential negative outcome more or less likely, imminent, or salient. This progression produces peaks and valleys in the suspense time course. Each excerpt was about 3 minutes long, with a total duration of 31 minutes 47 seconds.

### Procedure

Stimuli were presented using custom scripts written with the Psychtoolbox 3 software package for MATLAB. Participants viewed the 10 suspenseful film excerpts in a randomized order for each participant, with one functional run for each film excerpt, preceded and followed by 20.3 s of a fixation cross. Participants also completed a 306.6 s resting state scan. Participants were instructed to keep their eyes open and look at a centrally presented fixation cross but to otherwise think about anything.

### MRI Acquisition

Whole-brain MRI scans were acquired with a 3T Siemens MAGNETOM Trio scanner at the GSU/GT Center for Advanced Brain Imaging. First, a structural T1-weighted magnetization-prepared rapid-acquisition gradient echo (MPRAGE) scan was collected with 1 mm isotropic voxels, sagittal slice acquisition, field of view = 256 mm, TR = 2250 ms TE = 3.58 ms, flip angle = 9°, 176 slices, GRAPPA acceleration factor 2. Then, a functional resting state scan was collected, followed by a functional run for each of 10 movie excerpts in a randomized order. The functional runs used a T2*-weighted echo-planar imaging sequence with the following parameters: axial slice acquisition, interleaved, matrix = 64 × 64, field of view = 220 mm, TR = 700 ms, TE = 30 ms, flip angle = 90°, 22 slices, voxel size 3.4 × 3.4 × 3.1 mm, gap = 3.56 mm, slice acceleration factor = 2. At the beginning of each run, two additional volumes were discarded to account for T1 equilibration effects.

### Analysis

#### Identifying peaks and valleys of suspense

Dynamic suspense ratings for the stimulus films were taken from our other studies (Bezdek et al., 2015, 2017; Bezdek & Gerrig, 2017). An independent group of participants viewed the same ten suspenseful film excerpts while moving a joystick slider up and down to indicate how strongly the character was threatened over time. To identify the time course of when suspense was increasing and decreasing, the average of the derivative of joystick positions was computed across participants.

In the present study, we were interested in brain states near peaks and valleys of suspense. The change in suspense ratings, originally sampled at 10 Hz, were resampled to the rate of MRI acquisition and convolved with a canonical hemodynamic response function. We computed peaks and valleys using the following rules: the five highest maxima were recorded as peaks and the five lowest minima were recorded as valleys, and, to limit the influence of the pre- and post-movie periods on the suspense analysis, any peaks or valleys occurring within the first 10 or last 10 TRs were replaced with the next most extreme maximum or minimum.

#### Preprocessing MRI Data

A preprocessing pipeline was generated using the program afni_proc.py (version 4.38) of the AFNI software library (Cox, 1996). The pipeline included the following steps: de-spiking, aligning to the anatomical scan using the third volume of the first run, warping to MNI space, applying an eroded brain mask, using a custom MATLAB script to apply aCompCor (Behzadi et al., 2007) to regress nuisance signals from CSF and white matter regions of interest, applying spatial smoothing with a Gaussian kernel at full width at half-maximum (FWHM) = 6.0 to reduce noise, performing global signal regression, censoring TRs at which the Euclidean norm of the motion parameters was 0.2 or greater, and regressing demeaned and derivatives of head motion parameters.

#### Suspense GLM

A massive univariate general linear model was conducted on the functional MRI data, using the change in suspense ratings as a continuous parametric regressor. In addition to the suspense regressor of interest, the design matrix also included nuisance regressors for head motion in six directions, polynomial regressors for scanner drift, and the values for the following low-level visual features: average hue, saturation, luminance, and number of edges per frame, and the average magnitude of frame-to-frame dense optical flow. Individual beta weights for the suspense regressor were used for a group t-test. Monte Carlo simulations were generated using the program 3dClustSim to estimate the necessary cluster-size with a corrected family-wise error rate of 0.05. A *t*-threshold of 3.415 was chosen that limited the family-wise error rate to 0.05 with a minimum cluster size of 31 voxels.

## Results

### Brain States

In order to reduce computational complexity and leverage intrinsic functionally-correlated networks, the pre-processed BOLD time courses for the whole brain were parcellated into 14 functional networks using an atlas from Shirer and colleagues (2012). In that study, the authors identified the 14 networks by performing independent component analysis on group resting state data, and visually selecting and thresholding components that corresponded to intrinsic connectivity networks reported in prior research. The average BOLD response at each timepoint was averaged for each of the 14 networks. The time courses were then z-scored within each network.

*K*-means clustering was used to divide all functional run time-points into discrete states, and the number of states used for analysis was decided by the elbow method. Clustering was performed for each number of clusters from 2 to 12 and the proportion of sum of squared variance within-cluster over total variance was calculated for each cluster size. Then the second derivative of the resulting function was taken to determine at which *k*-value there was a local peak in the change in variance explained. This analysis determined that five clusters was a local peak for the curvature of variance explained and so the clusters derived from five states were used for all remaining analyses.

The centroids of the five brain states, derived from computing the arithmetic mean of activation for all time points assigned to each cluster, are shown in Figure 1a. State 1 was marked by increased activation in the posterior and anterior salience networks and visuospatial networks, with decreased activation in dorsal and ventral default-mode networks and the precuneus. State 2 was marked by widespread activation across all of the functional networks except the higher visual network. State 3 was marked by increased activation of the ventral default-mode network, with moderate activity in the precuneus and executive control networks and deactivation of the anterior salience network. State 4 was marked by activation of the dorsal default-mode network and moderate activation of executive control networks, with deactivation of sensory processing networks. Finally, State 5 was marked by widespread deactivation, particularly in executive control networks, and activation of visual and auditory networks.

**Figure 1.**
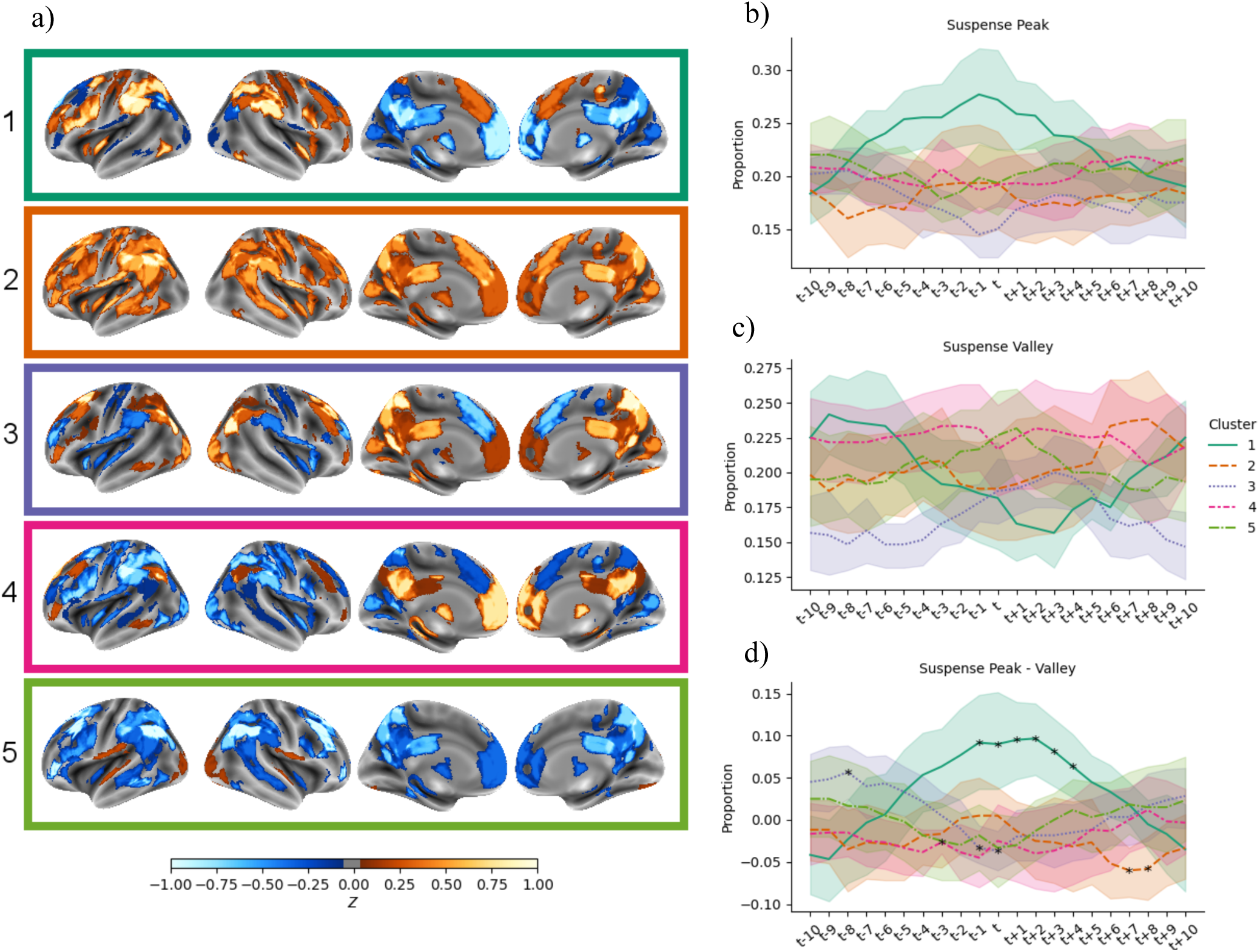
a) Centroids of the five clustered brain states, derived from averaging the activity in each network for all time points assigned to each cluster, then projecting to the FreeSurfer fsaverage inflated surface. The units of activation are *z*-scores of the preprocessed BOLD data. Proportion of brain states occurring in the time surrounding peaks (b) and valleys (c) of the suspense time course, and the difference in proportion of peaks minus valleys (d). A window of 21 volumes was used, centered at each local suspense peak or valley (time *t*) and each time step is 700 ms. Shaded areas represent 95-percent confidence intervals around each state mean. Points marked with asterisks in d) significantly differed from zero as tested with a sequential goodness of fit test.

Resting state volumes were assigned to clusters by computing the Euclidean distance between each resting volume and the centroid of each of the clusters derived from movie-viewing, then assigning the cluster with the smallest distance.

### Likelihood of Brain States at Suspense Peaks and Valleys

Using the identified peaks and valleys of suspense, we computed the frequency of occurrence of each brain state at each whole-brain volume (one for each TR) in a window surrounding the volume of each peak or valley, including 10 volumes before and 10 volumes after. Then we computed the relative frequency between peaks and valleys to identify states that were more or less likely to occur at each TR of the window around peaks than valleys. Finally, a sequential goodness-of-fit test was used to correct for multiple comparisons when judging if any of the states were significantly different from zero across the window (Carvajal-Rodríguez et al., 2009).

Four of the five states included time points that were significantly more or less likely at peaks than valleys (see Figure 1b-d). State 1, marked by increases in the salience network and decreases in the default-mode network, occurred significantly more frequently at suspense peaks than valleys, from time *t* – 1 to time *t* + 4 (−0.7 to 2.8 s). State 2, marked by widespread activation, was significantly more likely at valleys than peaks at time *t* +7 and *t* + 8 (4.9 to 5.6 s). State 3, marked by increased activation in the ventral default-mode network and anterior salience network deactivation, occurred more frequently at peaks than valleys at time *t* – 8 (−5.6 s), then more frequently at valleys than peaks at times *t* – 1 and *t* (−0.7 to 0 s). Finally, State 4, marked by dorsal default-mode network activation, was more common at valleys than peaks at time *t* – 3 (− 1.4 s).

### Transition Probabilities Between States

We calculated the proportions of transitions between states from one time point to the next, for both film viewing and resting state data. Film viewing and resting produced similar patterns of state transitions (see Figure S1a and S1b). Computing a paired-samples *t*-test by subject for the 11 subjects with both film viewing and resting state data did not produce any significant differences in state transitions that survived correction for multiple comparisons. We also computed transition proportions separately at peaks and valleys by taking the state at *t* + 2, where the largest difference between peaks and valleys was observed, then recording the next state that occurred up to 15 TRs forward. Figure S1c and S1d show the average transition proportions at peaks and valleys and Figure S1e shows the average difference between peak and valley transition proportions. When computing paired-samples *t*-tests by subject, there were no significant differences between peak and valley transitions that survived correcting for multiple comparisons.

### Univariate Activation in Response to Changes in Suspense

Figure 2 shows univariate increases and decreases in activation as a function of suspense. Moments of suspense evoked significant increases in activity in bilateral occipitotemporal cortex. Significant increases in activation were also found in the right temporoparietal junction and the right middle frontal gyrus. Decreased activation was found in the posterior cingulate cortex and medial occipital cortex. These results replicate previous studies using this technique (Bezdek et al., 2015, 2017), that reported suspense related decreases in activation in the default-mode network and increases in activation in the ventral attention network and visual association cortices.

**Figure 2.**
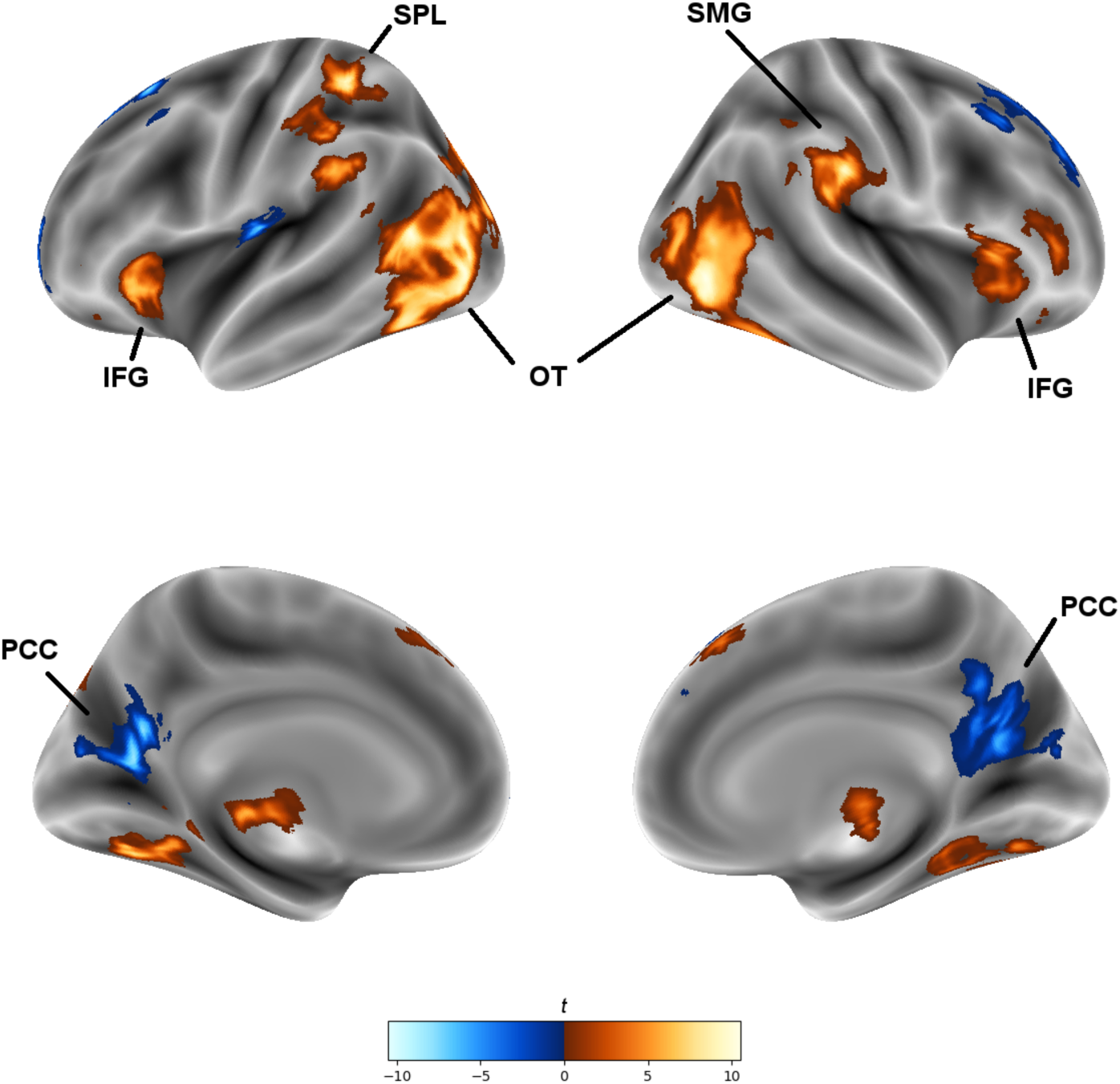
Univariate response to changes in narrative suspense. Suspense produced univariate increases in lateral occipito-temporal cortex (OT), right supramarginal gyrus (SMG), bilateral inferior frontal gyrus (IFG), and left superior parietal lobule (SPL). Suspense produced univariate decreases in activity in the posterior cingulate cortex (PCC). Clusters of activation that survived correction to a family-wise error rate of 0.05 are shown projected on the inflated FreeSurfer fsaverage surface.

## Discussion

We used a fast multiband acquisition sequence to investigate the brain network state configurations during naturalistic suspenseful film viewing. We identified five brain states of activity patterns during suspenseful film viewing. These states bear resemblance to those found in other studies using resting state and task data. In particular, Smith et al. (2018) found states of widespread activation, deactivation, and strong default-mode activation when clustering resting state data. Chen et al. (2015) reported similar spatial patterns in dominant co-activation patterns derived from resting state scans and working-memory task scans. We did not find significant differences in the likelihood of state transitions between movie viewing and rest, or between suspense peaks and valleys. Comparing the likelihood of brain states at suspense peaks and valleys revealed the fluidity of brain states at these critical moments within the narrative. In particular, suspense peaks were associated with a brain state marked by high activity in the salience network. Interestingly, the dominance of this state began 0.7 seconds before the suspense peak, which is especially early when taking into account the lag in hemodynamic response. This suggests that viewers were already in a state of heightened engagement as the increase in suspense was peaking, perhaps due to preceding narrative information or viewers’ predictive processing. States 3 and 4, which split the dorsal and ventral components of the default-mode network, were each more prevalent at valleys than peaks at times within the window. State 2 (characterized by widespread activation) appeared significantly more frequently about 5 seconds after a valley than after a peak, perhaps due to suspense peaks evoking the more focused network recruitment present in the other states rather than the diffuse activation of State 2.

The pattern of univariate responses to suspense replicates previous findings on brain activation evoked by suspenseful film moments (Bezdek et al., 2015, 2017). In particular, the right-lateralized activity in the temporo-parietal junction and the middle temporal gyrus overlaps with regions identified in previous work as the ventral attention network, a network recruited to direct attention to arousing stimuli in the environment (Corbetta & Shulman, 2002). The deactivation in medial occipital cortex mirrors a previous finding that narrative suspense produces deactivation in regions of occipital cortex that process the visual periphery when suspenseful films are presented at fixation with continuous peripheral stimulation. These results support the theory that peaks in suspense reorient neural processing to the narrative and suppress processing of peripheral visual stimuli, confirming the reliability of univariate brain effects of suspense in a novel sample and with a fast multiband acquisition sequence.

Comparing the univariate and brain state results provides a fuller picture of the dynamics of narrative suspense. While the univariate response to suspense shows decreased activation in the PCC, using the brain state analysis reveals three different states (2, 3, and 4) with increased PCC activation that are more prevalent in the time surrounding valleys than peaks. Though these states share recruitment of PCC, they differ in the pattern of activation of other networks across the rest of cortex. In contrast, the increased activation in the ventral attention network is associated with a single state (1) that is prevalent at suspense peaks. Together, these results reveal that suspense peaks evoke a common brain state of engagement, while suspense valleys evoke more variable brain states.

If suspenseful moments evoke attentional narrowing and reorienting to narrative stimuli, what are the consequences for what follows? One theory is that suspense evokes viewer participation on the behalf of the characters in danger. Gerrig and Bernardo (1994) reported evidence of readers engaging in problem-solving when a protagonist faced a suspenseful situation. When items that the protagonist could use to escape were removed from him, readers rated the story as more suspenseful. When viewers of suspenseful movie clips were asked to speak their thoughts aloud, they also made responses that indicated problem-solving on behalf of characters (Bezdek et al., 2013). Brain states that reflect mental participation, simulation, and problem-solving may appear at other times during the viewing of suspenseful films. State 3, with its combination of executive control and default-mode network activation, is one candidate state for capturing the mental participation of engaged viewers. A potentially fruitful avenue for future research is to more directly probe the downstream effects of suspense on viewers’ mental participation and memory.

The use of functional networks to define brain state clusters has both advantages and drawbacks. Averaging voxel-level BOLD responses to a smaller number of network activations reduces the computational load and may reduce noise present at a voxel level. However, caution must be exercised in inferring network function from observed activation. Further, it is possible that dynamic network reorganization may occur during these tasks, producing new patterns of communication between network sub-regions. Examining the dynamics of network configurations during naturalistic tasks is another area ripe for further investigation.

To be consistent with other related literature (e.g., Fox et al., 2009; Li et al., 2019; Power et al., 2011), we included global signal regression (GSR) in our pipeline. There is disagreement in the literature over the use of GSR, because in addition to noise there may also be meaningful information in the global signal (Murphy & Fox, 2017). Anti-correlations between regions may increase after performing GSR. However, using a similar clustering method to identify brain states in resting state fMRI data, Smith and colleagues (2018) reported commonalities in the spatial patterns of brain states derived with and without performing GSR, though the states of widespread activation and deactivation were absent when GSR was included, unlike in our study. Due to the unknown and controversial tradeoffs in performing GSR, we recommend caution in interpreting whether the contrasting relationship between salience and default-mode networks that we see in States 1 and 3 reflects a true anti-correlation or a difference in degrees of activation. Although, GSR is less likely to have a large effect on our other results.

In conclusion, we find that narrative suspense produces different patterns of brain states at peaks and valleys of the suspense time course. This finding is compatible with theories of suspense as an attentional spotlight that triggers subsequent cognitive engagement of viewers with the narrative world. Our results demonstrate the power of multiband fMRI clustering analyses to investigate moment-by-moment changes in brain states during naturalistic tasks at a sub-second time scale.

