## Supplementary Materials for "Dynamic Brain Network States during Suspenseful Film Viewing"

### Supplementary Material

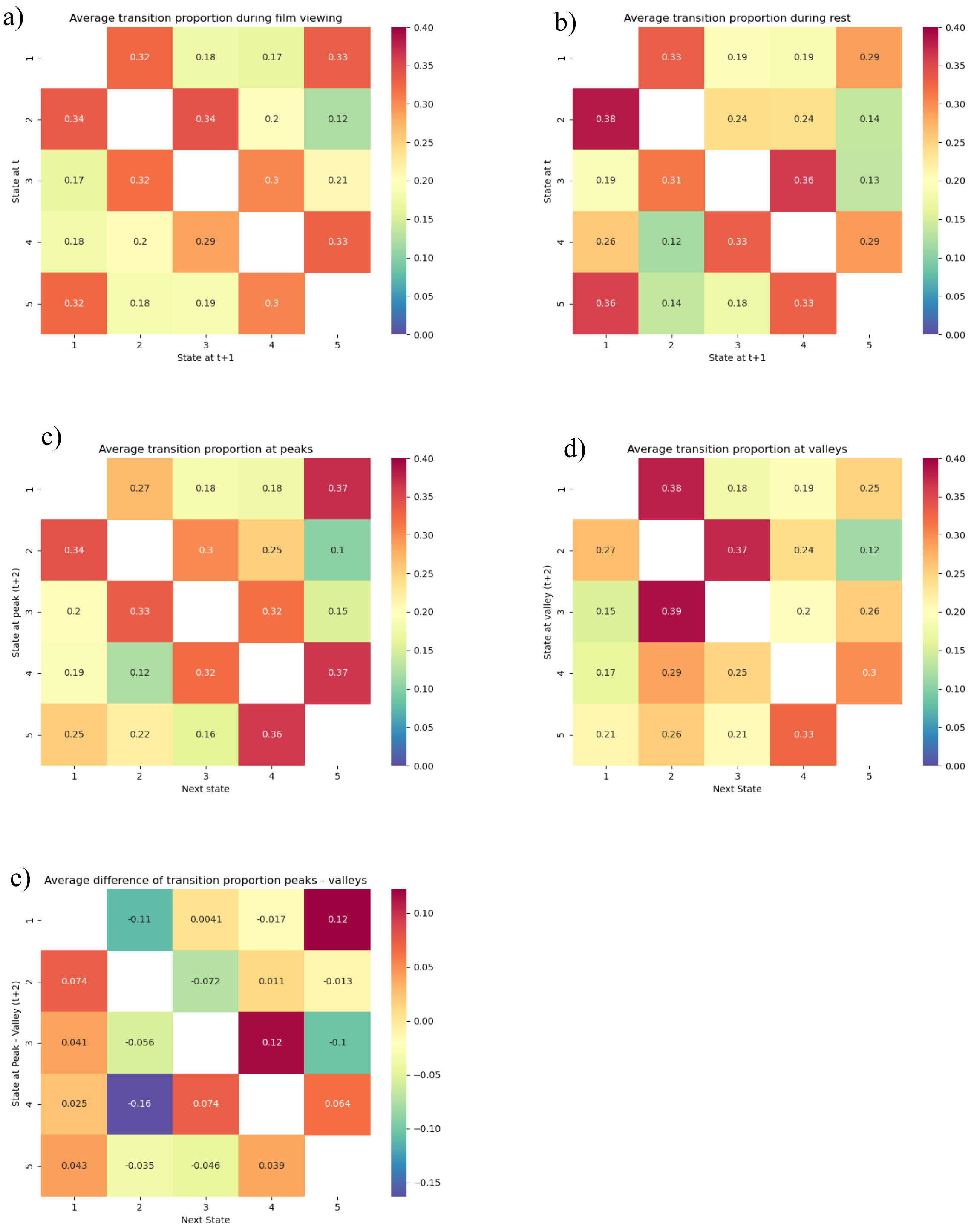


Supplementary Figure 1. Average state transition proportions during film viewing (a) and during resting state scans (b). State transition probabilities were generally similar for both film-viewing and rest despite the differences between the two tasks. Transitions at suspense peaks (c) and suspense valleys (d), starting at two TRs after a peak or valley and measuring the next state within 15 TRs, showed similar transition probabilities, and differences between peak and valley transitions (e) was not significant across participants after correcting for multiple comparisons.

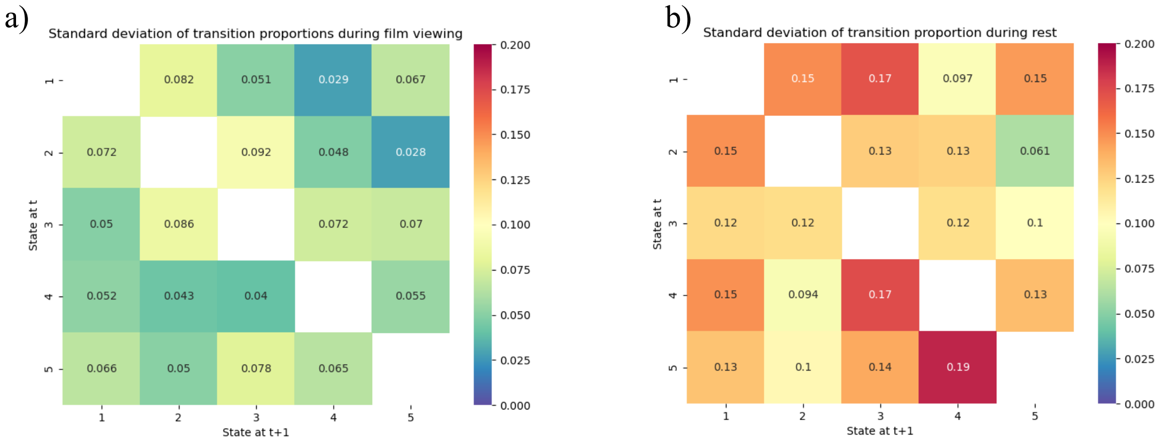


Supplementary Figure 2. Standard deviation of state transitions during (a) film viewing and (b) resting state. Resting state transitions are calculated using a much smaller set of observations than the film viewing transitions, resulting in larger standard deviations overall.
